# Structure of a SARS-CoV-2 spike S2 subunit in a pre-fusion, open conformation

**DOI:** 10.1101/2023.12.14.571764

**Authors:** Eduardo Olmedillas, Roshan R. Rajamanickam, Ruben Diaz Avalos, Fernanda A. Sosa, Michelle A. Zandonatti, Stephanie S. Harkins, Sujan Shresta, Kathryn M. Hastie, Erica Ollmann Saphire

## Abstract

The 800 million human infections with SARS-CoV-2 and the likely emergence of new variants and additional coronaviruses necessitate a better understanding of the essential spike glycoprotein and the development of immunogens that foster broader and more durable immunity. The S2 fusion subunit is more conserved in sequence, is essential to function, and would be a desirable immunogen to boost broadly reactive antibodies. It is, however, unstable in structure and in its wild-type form, cannot be expressed alone without irreversible collapse into a six-helix bundle. In addition to the irreversible conformational changes of fusion, biophysical measurements indicate that spike also undergoes a reversible breathing action. However, spike in an open, “breathing” conformation has not yet been visualized at high resolution. Here we describe an S2-only antigen, engineered to remain in its relevant, pre-fusion viral surface conformation in the absence of S1. We also describe a panel of natural human antibodies specific for S2 from vaccinated and convalescent individuals. One of these mAbs, from a convalescent individual, afforded a high-resolution cryo-EM structure of the prefusion S2. The structure reveals a complex captured in an “open” conformation with greater stabilizing intermolecular interactions at the base and a repositioned fusion peptide. Together, this work provides an antigen for advancement of next-generation “booster” immunogens and illuminates the likely breathing adjustments of the coronavirus spike.

## INTRODUCTION

Multiple families of viruses that cause severe human disease have an essential surface fusion protein. This protein, whether termed *env* in retroviruses, GP in filoviruses, GPC in arenaviruses, hemagglutinin in influenza viruses, or spike in coronaviruses, is the essential piece of viral machinery that drives the virus into the cell. These critical proteins are targeted by neutralizing antibodies (nAbs) and are a major component of vaccines. Both antibodies and vaccines are challenged, however, by mutagenic substitutions that escape antibody recognition. Developing vaccines that target conserved, unchanging parts of the fusion protein is key for long-lasting, broadly active immune protection.

For many of these viruses, the fusion protein contains two subunits: one which mediates membrane fusion and the other which mediates initial receptor binding. Typically, the receptor-binding subunit (variously termed gp120, HA1, GP1, S1, etc.) is more subject to selective pressure and mutation, while the fusion subunit (variously termed gp41, HA2, GP2, S2, etc.) remains more conserved in sequence. The two subunits are formed by enzymatic cleavage of a polyprotein precursor in the producer cell, and although cleaved apart, remain associated on the viral surface. As long as they are associated, one (the receptor-binding subunit) serves as a clamp on the conformation of the other (the fusion subunit). On the surface of the virus, the fusion subunit exists in a metastable, initial, pre-fusion conformation, interacting with or intertwined with the receptor-binding subunit. Upon attachment to the receptor and often, exposure to low pH, the receptor-binding subunit releases its clamp and the fusion subunit irreversibly rearranges into a more stable, six-helix bundle structure called the “post-fusion conformation”. This energetically favorable conformational rearrangement can thwart the neutralizing antibody response: multiple neutralizing antibodies have been discovered that target only the metastable prefusion conformation and are unable to recognize the post-fusion, six-helix bundle conformation (*1–12*). The greater inherent stability of the post-fusion six-helix bundle means that without its clamping receptor-binding subunit, the fusion protein, when expressed alone, adopts only the post-fusion six-helix bundle conformation. The metastability of the fusion subunit has challenged development of immunogens or research reagents that constitute the prefusion S2 (or gp41) alone.

Difficulties in making fusion subunits alone in their pre-fusion conformation is a loss for immunogen development, as the fusion subunit is more conserved than the receptor-binding subunit. Between SARS-CoV-2 and SARS-CoV, for example, the S2 fusion subunits are 90% conserved while the S1 N-terminal domain and receptor binding domains are 50% and 76% conserved, respectively (*13*). Targeting S2-reactive immune responses offers an opportunity to build cross-reactive protection bridging from previous coronavirus infections, and extending to coronaviruses yet to emerge. As an example, convalescent sera from patients infected with SARS-CoV or SARS-CoV-2, who were unlikely to have ever been exposed to MERS, nonetheless include antibodies that react with or neutralize MERS (*14, 15*); and immunization with S2-based constructs elicited a broadly cross-reactive IgG antibody response that recognized the spike proteins of not only SARS-CoV-2 variants, but also SARS-CoV-1 and the four endemic human coronaviruses (*16*). Meanwhile, other individuals, naïve to SARS-CoV-2, had immune responses that reacted with regions of its S2, presumably from prior common cold CoV infections (*17*). Several known S2-reactive antibodies are neutralizing and protective against infection and pathology *in vivo* (*18–20*). “Boosting” with the conserved fusion subunit after a whole-envelope spike would be an attractive strategy for building broader immunity (*21*), but unfortunately, the wild-type fusion subunit for these viruses can not be expressed alone in the correct pre-fusion conformation.

In addition to the dramatic and irreversible pre-fusion to post-fusion refolding event, the viral surface glycoproteins are also thought to exhibit reversible, conformational “breathing” motions (*22, 23*). A low-resolution structure of the respiratory syncytial virus (RSV) F trimer, for example, suggests that F protein exists in both open and closed conformations (*24*). Breathing motions have been described for hantaviruses (*25*), flaviviruses (*25, 26*) and retroviruses. Motion in the envelope (Env) protein of HIV-1 has been observed in both soluble and virion-surface forms (*27*), (*28*), and is thought to be required for binding of the HIV-1 receptor and coreceptor. Antibodies that cause *Env* trimer dissociation have been described and may depend on the same breathing motions required for receptor binding. For SARS-CoV-2, molecular simulations of spike protein also suggest a dynamic prefusion state, including opening of the monomers from the trimeric stem interface (*29*). This motion could enhance accessibility of receptor-binding domains and expose the conserved trimer interface for recognition by antibody (*30*). Although supported by molecular dynamics and biochemical and biophysical measurements, the structure of an open state of SARS-CoV-2 has not yet been determined.

In this work, we have engineered the fusion-mediating S2 subunit of SARS-CoV-2, to stably remain in its prefusion conformation alone, in the absence of the receptor-binding S1 clamp. We provide evidence from electron microscopy that the engineered S2 indeed stably remains in its prefusion conformation. We also describe here competition mapping of novel human mAbs against S2 from both convalescent and vaccinated individuals, studies which were facilitated by the new existence of a pre-fusion S2-only antigen. One of these mAbs allowed the determination of a cryo-EM structure at 2.9 Å of the prefusion S2. Notably, this structure illustrates a high-resolution view of S2 in an open conformation that reflects prior biophysical predictions. We find that human mAbs from both convalescent (not yet vaccinated) and vaccinated (but not yet infected) individuals react with an upper, inner surface of the open S2, an epitope that is accessible in a “breathing” open S2, but masked in a closed S2, suggesting that the breathing motion of the full-length spike required to expose this epitope happens in both natural infection and in the context of vaccination. Notably, the open conformation of S2 reveals a greater number of intramonomer stabilizing interactions in the trimer base as well as an alternate position and conformation of the fusion peptide.

This work thus not only provides a novel pre-fusion S2-only immunogen and research tool but also a high-resolution view of a conformational state of spike relevant for antibody recognition and the immune response. The over 800 million human SARS-CoV-2 infections, coupled with the likely emergence of new variants and new coronaviruses, necessitates development of additional vaccine strategies featuring conserved sites such as S2 that foster broad and durable immunity.

## RESULTS

### Design of the SARS-CoV-2 stem domain in the prefusion conformation

Introduction of disulfide bonds can substantially increase the stability of viral glycoproteins, as has been demonstrated for HIV, LASV and RSV (*4, 5, 7, 21, 31*). We previously described the development of an engineered SARS-Cov-2 spike protein, termed S_VFLIP_, that contains five (V) proline substitutions (one subtracted from Hexapro to restore a native, stabilizing salt bridge between K986 and D748 (*32*)), a flexible linker (FL) in place of the native furin cleavage site, as well as a novel inter-protomer (IP) disulfide bond between residues Y707C-T883C, located in the interior of the S2 stem which maintains spike in its trimeric organization without an exogenous trimerization domain (*33, 34*). The S1 region of this construct, and the FL from VFLIP, were genetically removed to create a S_VFLIP_ S2-only construct (residues 691-1208) to serve as the basis for further engineering.

To stabilize the prefusion conformation of S2 in the absence of S1, we used the Disulphide by Design server (*35*) to computationally predict optimal disulfide bonds in the VFLIP-S2 ectodomain. Of the 31 pairs identified, we found that ten were likely to form the stabilizing disulfide bond in the pre-fusion (PDB: 6XKL), but not the post-fusion conformation (PDB: 7E9T) (*36–38*).

Each of these ten intra-monomeric disulfide bonds was engineered into the S2 subunit bearing VFLIP substitutions. To assess the viability of these constructs as potential vaccine antigens, we comprehensively examined the feasibility of high-yield expression for each of the ten constructs in ExpiCHO cells, as well as conformation, protein aggregation and stability. We also expressed the wild-type S2 ectodomain (residues 691-1208), which naturally refolds into the post-fusion conformation, as a control. Two constructs, DS2 and DS3, had minimal-to-no expression and were not characterized further. The remaining eight constructs, however, yielded a sufficient amount of protein to evaluate by size-exclusion chromatography (SEC). Seven of the eight constructs eluted as a major peak at 14 mL, with only minor shoulder and aggregation peaks. The eighth, DS1, eluted earlier, at 13 mL. Of note, four of the engineered constructs (DS1, DS5, DS6, and DS9) yielded more than more than 5 mg purified, monodisperse protein from a 50 mL culture of transiently transfected ExpiCHO cells (extrapolated yield: >100 mg/L) (Fig.1B and Suppl. Table T1). Non-reducing SDS-PAGE showed that all four proteins retain a covalently-linked trimeric state with efficient disulfide bond formation, as evidenced by a band at the expected molecular weight of 200 kDa and a lack of detectable low molecular weight species (Fig. 1D). We did not observe higher molecular weight species that might indicate aberrant disulfide bond formation. Meanwhile, under reducing conditions, all constructs exhibited the monomeric molecular weight of the wild-type S2 (∼80 kDa), suggesting that trimerization was anchored as expected by the Y707C-T883C disulfide bond.

**Figure 1.**
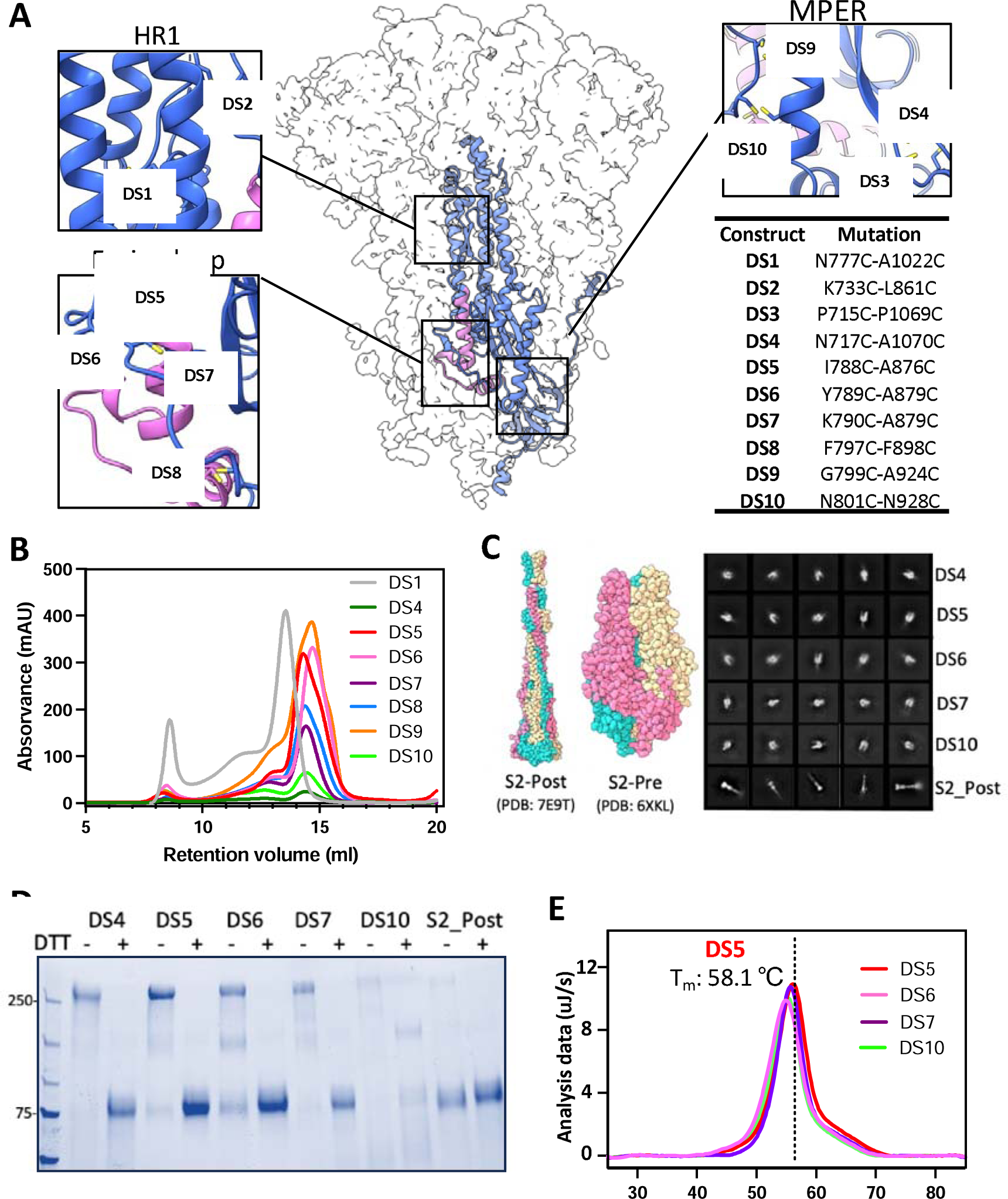
Characterization of covalently-linked SARS-CoV-2 S2 variants. **(A)** Side view of the trimeric SARS-CoV-2 spike (silhouette) with a schematic representation of a monomer of S2 (PDB:6XKL) in ribbon diagram. The fusion peptide (FP) is colored in magenta. Each inset corresponds to the location of the computationally beneficial introduction of cysteines to form disulfide bonds. The position of each substitution is shown in the table. **(B)** Size-exclusion chromatography (SEC) traces comparing S2 purity post-purification. **(C)** Atomic diagram of the post- and pre-fusion conformations with each monomer colored differently. Representative negative-stain electron 2D classes of the S2-prefusion candidates DS4, DS5, DS6, DS7, and DS10, showing homogenous well-formed trimer populations, and S2 post-fusion construct from wild-type S2. **(D)** Non-reducing/Reducing SDS-PAGE of constructs shown in (C). **(E)** Differential Scanning Calorimetry (DSC) analysis of spike thermostability.

Negative stain electron microscopy (ns-EM) showed that constructs DS4, DS5, DS6, DS7 and DS10 adopt the expected ‘pear’ shape of S2 in its prefusion state. In contrast, wild-type S2 adopted only the characteristic stick shape of the post-fusion conformation (2D classes, Fig 1C, Table S1). The remaining constructs were highly heterogeneous and did not yield any clear 2D classes. We next evaluated the thermostability of DS4, DS5, DS6, DS7 and DS10 by Differential Scanning Calorimetry (DSC). Of these, DS5 exhibited the highest thermostability with a T_m_ of 58.1±0.1 °C, compared to the lowest T_m_, 54.8±0.1 °C, of DS6 (Fig. 1E).

We moved forward with the best engineered S2 construct, S2-DS5, which stably presents only the pre-fusion conformation, and exhibits one of the highest expression yields (∼150 mg/L; similar to full-length S_VFLIP_ ectodomain) and the greatest thermostability, suggesting its utility as both a potential immunogen and a research tool to elicit and map pre-fusion S2-specific antibodies.

### Binding kinetics and antigenic communities of anti-S2 antibodies isolated from human subjects

Most antibody epitope mapping efforts have focused on the S1 subunit and its receptor-binding domain. Comparatively fewer antibodies are known, and fewer epitopes mapped, in the S2 subunit. We previously isolated and characterized a panel of antibodies that target the spike protein from convalescent (unpublished) and vaccinated (*39*) subjects. Approximately one-fourth react to regions outside the N-terminal and receptor-binding domains of S1, suggesting possible recognition of the S2 subunit. We first used high-throughput SPR (Carterra LSA) to map the conformational specificity using S_VFLIP_-D614G, S2-DS5, and post-fusion S2 and evaluated cross-reactivity of these anti-S2 antibodies to S2-DS5-XBB1.5, S_VFLIP_-XBB1.1, and S_VFLIP_-SARS-CoV-1. Of the 48 mAbs tested, two recognized full-length D614G Spike (S1+S2 domains) only, with no recognition of pre-fusion S2-DS5 or post-fusion S2 alone, suggesting that their epitopes involve S1 in some way. Two mAbs were specific for post-fusion S2, with very weak or non-existent binding to full spike or pre-fusion S2-DS5. The remaining 44 mAbs, however, recognized both full-length Spike and the pre-fusion stabilized S2. One mAb (3B11) recognized S2-DS5 but not post-fusion S2, suggesting this mAb may bind to an epitope presented only in the prefusion conformation of S2. Aside from this single mAb, other Spike/DS5-S2-reactive mAbs also bound to post-fusion S2, and curiously, often did so with higher affinity. We also found that 40 mAbs bound to XBB1.1 and SARS-CoV-1 spikes (Fig. 2A).

**Figure 2.**
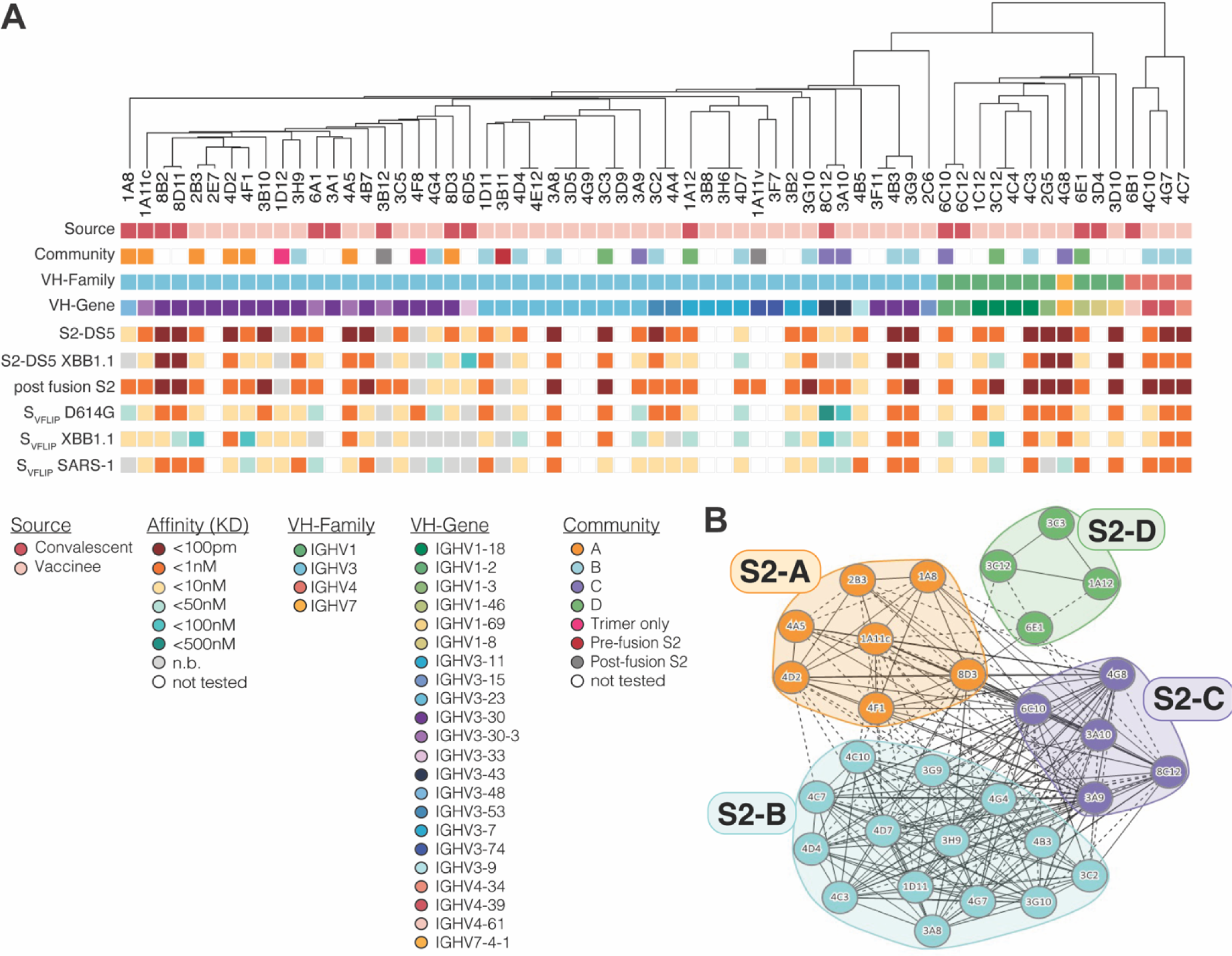
The kinetics and antigenic landscape of S2-directed mAbs. **(A)** A phylogenetic tree based on the heavy chain of each mAb is shown, with the indicated features of each mAb displayed below. **(B)** Pair-wise competition analysis was used to map the epitope community for a sub-set of S2-DS5 reactive mAbs. Circles indicate individual mAbs with larger clusters grouped together and colored by community. Solid lines connecting mAbs indicate that the two antibodies compete bi-directionally; dotted lines indicate asymmetric competition.

We next sorted 30 of the S2-DS5 reactive mAbs into epitope communities. S2-DS5 was preincubated with >10x the K_D_ of each mAb and complexes injected over the same array used for kinetic studies. This panel of mAbs were sorted into four communities (S2-A to S2-D) (Fig. 2B). Interestingly, the S2-A community, which contains mAbs from both convalescent and vaccinated subjects, may represent a convergent solution to this particular antigenic region, as all but two members originate from the 3-30 VH germline.

### Structural characterization of an engineered S2 domain

Representative nsEM 2D classes of the antigen-only S2-DS5 structure demonstrate that it adopts the pre-fusion conformation (Figure 1D). To add molecular mass and a fiducial to improve cryoEM resolution, we made complexes of S2-DS5 with an antibody from each of four representative communities, and evaluated them by nsEM (Suppl. Fig S1). One complex, of Fab 6C10 bound to S2-DS5, yielded a good distribution of particles and orientations, including top-down and side views, which was suitable for high-resolution 3D structure determination (Suppl. Fig. 1). Antibody 6C10 was identified in a convalescent individual, from PBMCs donated early in the pandemic prior to the availability of vaccines. Two datasets from two batches of the complex were independently collected (Suppl. Fig. 2). A final set of 407,314 particles of the complex led to a 2.9 Å reconstruction, with C1 symmetry and no masks applied (Fig 3A). Three copies of the 6C10 Fab fragment are bound to the prefusion S2 trimer (Suppl Table 2).

**Figure 3.**
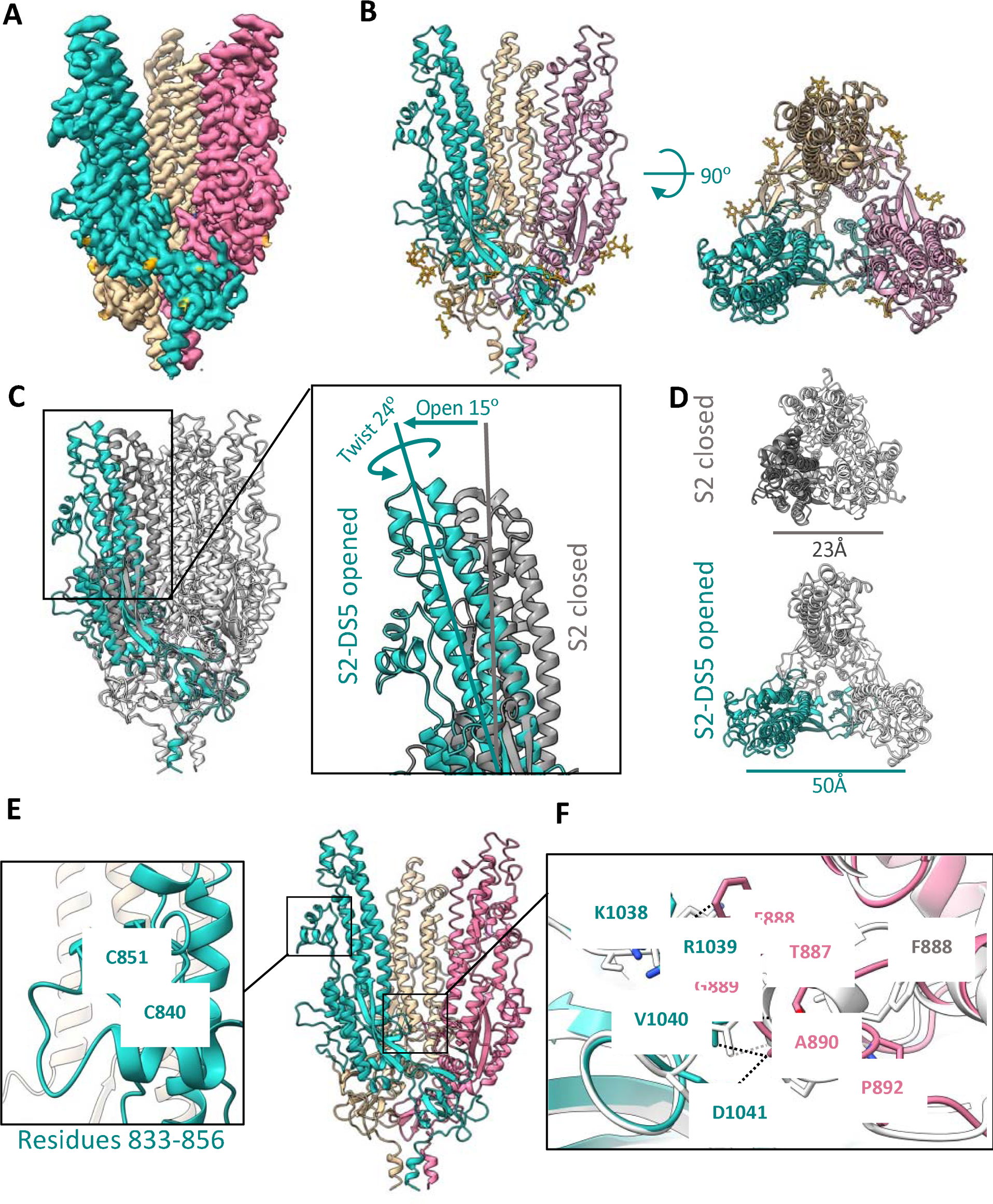
Structure of the prefusion SARS-CoV-2 S2 subunit. **(A)** Side view of the cryo-EM density map of the trimeric SARS-CoV-2 S2 subdomain. Each protomer is colored differently, and glycan densities are colored orange. **B)** Side (left) and top (right) views of the S2-DS5 molecular model, colored as in (A). **(C) and (D)** Alignment of S2-DS5 with the S2 from the full-length stabilized Hexapro (PDB:6XKL). (C) shows a side view of both trimers with one protomer colored in turquoise (S2-DS5) and grey (Hexapro). Zoomed view of the ‘open state’ of S2-DS5 by an opening and twisting motion. (D) shows the top views with the differences in distances between the P987 of two neighboring protomers. **(E)** S2-DS5 molecular model zooming the not previously solved loop 833-856, and **(F)** the novel inter-protomeric interaction between the fusion peptide loop 882-898 (magenta) and the loop 1033-1037 (turquoise), in comparison with S2 in the close state from HXP (structure showed in grey) Contacts are shown in black dash lines for S2-DS5 and grey for HP.

In the map, residues 706-1149 of S2 can be modeled (Fig. 3B). As in pre-fusion structures of the complete spike, the C-terminal peptide (residues 1150-1208) is not visible. We do, however, observe density for each of 6 N-linked glycans attached to each S2 monomer (18 per trimer) located in positions N709, N717, N801, N1074, N1098, and N1134. The newly engineered I788C-A876C intra-monomer disulfide, which holds S2 in the pre-fusion state, and the S_VFLIP_-derived Y707C-T883C inter-monomer disulfide, which maintains the trimer in the absence of an exogenous domain, are also both clearly ordered in the cryo-EM maps. We further observe clear density for a loop comprising residues 833-856 located in the equatorial part of the full-length spike, under the CD1 domain (Fig. 3E and Suppl Figs. S3 and S4). This loop has been poorly ordered in all pre-fusion structures of spike thus far (40–45).

### The breathing conformation

Previous biophysical experiments demonstrated that S2 adopts closed and open states in the context of complete spike, whether wild-type or moderately stabilized by two prolines (*30*). However, all previous structures are of a similar closed conformation, in which a tightly trimerized S2 is capped by S1. The high-resolution structure of the open conformation remains unknown.

Comparison of the S2 in the S2-DS5/6C10_Fab complex presented here to the S2 in previous spike structures, shows that this complex has captured a more open conformation. The difference in structure between this open and previous closed states is essentially a rigid-body, screw-opening motion from the bottom membrane-proximal region (MPER) toward the upper reaches of each S2 protomer. This motion opens the monomers from the vertical axis by 10° degrees with a concomitant rotation of each protomer apex counter-anticlockwise 26° degrees (Fig. 3C and Suppl. movies 1 and 2). In the open state, the apex of each protomer (defined by the position of Pro 987) is 50 Å; in the closed state, the distance between each protomer’s Pro 987 is 23 Å (Fig. 3D). Analysis of S2 trimers using the PDBePISA server reported a total solvent-accessible surface of 22,300 Å2 in the closed state and 23,700 Å2 in the open state. In the closed conformation, the S2 trimer buries an average of 2,230 Å2 in each interprotomeric interface with a predicted ΔG =D-20.5DkcalDmol−1, whereas the open S2 trimer buries 1,260 Å2, with a predicted ΔG =D-20.1DkcalDmol−1.

The protomers themselves only differ between closed and open states by 0.76 Å r.m.s.d.; hence the motion for the most part is rigid body with each protomer top rotating open relative to the others. At the base, where the three protomers meet, the β-sheet-containing membrane-proximal region (MPER) (residues 1070-1138) from each monomer mediates interactions with its neighbor. These interactions, together with a short α-helix before the N-termini of HR2 (residues 686-705), confer most of the quaternary interactions visualized in the cryo-EM S2-DS5 map.

Interestingly, the open conformation of S2 appears to stabilize interprotomer interactions formed by the wild-type residues in the S2 base. This is perhaps paradoxical that an open conformation induces more stable interactions, so restated, the opening action of the upper part of S2 acts to stabilize and anchor the *base* where the protomers connect. In full, closed spike, the upper parts of S2 interact with S1. Relaxing the top into an open conformation may favor stabilization in the base; as the rigid body expansion of the top occurs with a concomitant compression of the base, satisfied by formation of chemically favorable interactions. The resulting greater number of stabilizing interactions in the open conformation is most visible in loop 885-897 in the base, which corresponds to the recently described fusion peptide of the spike (*38*). In closed structures, whether formed by S-2P or Hexapro spikes, residues 886-891 form a short α-helix, residues Phe888 and Gly880 of the same protomer are in close contact with each other, and Gly889 and Ala890 interact with Lys1045, Gly1046, and Tyr1047 from the 1037-1047 loop of the adjacent protomer, with a predicted ΔG = −3.3DkcalDmol−1 for the closed-state interactions (Fig 3F). In the open structure, there is an elongation of the 885-897 fusion peptide loop towards the 1037-1047 loop of the neighboring protomer. Residues Phe888, Gly889 and Ala890 are instead in closer contact to 1038-1041 (Lys-Arg-Val-Asp) (Fig. 3F), with a predicted ΔG = −4.0DkcalDmol−1 for the open-state interactions. Note that the structural differences in the base are related to the open conformation and not engineering or antibody binding, as all of these contacts are made by wild-type residues that are distant from the DS5 disulfide and distant from the Fab 6C10 footprint. Thus, this newly visualized interaction at the S2 trimeric base connection point may be important to maintaining the integrity of the trimer in the splayed-open or breathing conformation.

### Structural definition of the 6C10 epitope

Cryo-EM analysis of 6C10 Fab bound to S2-DS5 shows stoichiometric symmetry, in which one copy of the 6C10 Fab is stably bound to each protomer of S2-DS5 (Fig. 4A-B). The heavy- and light-chain complementarity determining regions (CDRs) bind ∼1,290 Å2 of the molecular surface on each S2 protomer, with the majority of the interface area, ∼87%, contributed by the heavy chain. 6C10 Fab approaches the S2-DS5 protomer from the side and bridges a novel quaternary epitope formed between the regions 733-772 and 844-862 of the apex of the S2 domain. Those two regions have no defined function yet, other than flanking the S2’ cleavage site (808-820).

**Figure 4.**
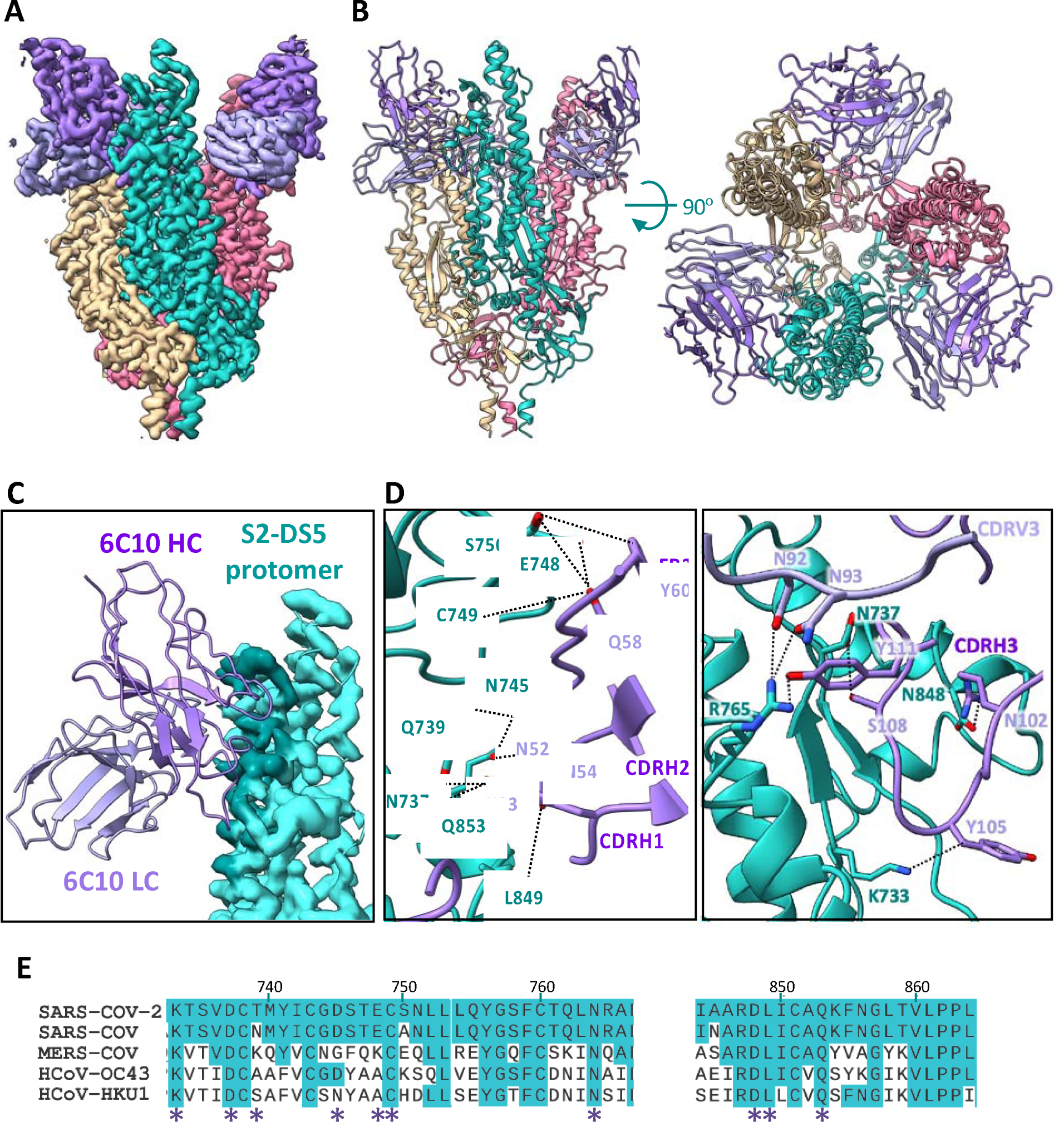
Cryo-EM analysis of the 6C10Fab/S2-DS5 interaction and relative conservation of epitope contact residues. **(A)** Side view of the cryo-EM density map of S2-DS5 ectodomain in complex with Fab 6C10. S2-DS5 is colored as in Figure 3, and the 6C10 Fab density is colored dark purple (heavy chain) and light purple (light chain). **(B)** Side (left) and top (right) views of the 6C10Fab/S2-DS5 molecular model complex, colored as in (A). **(C)** Zoom of the footprint of one 6C10 Fab to one S2-DS5 protomer. The 6C10 Fab is shown as ribbon representation and the S2-DS5 as the cryo-EM density. The footprint is colored darker than the rest of the S2. **(D)** S2-DS5 interactions with 6C10. The left panel shows a close-up view of interactions made by 6C10 CDRH1, H2, and framework region 2. Right panel shows the 6C10 CDRH3 interactions and the interaction of the 6C10 light chain (N92 and N93) with the residue R765 of S2-DS5. **(E)** Sequence alignment of the 6C10 conformational epitope (spike region 733-765 and loop 833-856) from the five human-infective beta-coronaviruses. Similarity in conservation with SARS-CoV-2 is shown in turquoise, and the interaction positions with the Fab are marked with a purple asterisk.

The majority of the 6C10 paratope is contributed by CDRH3. The 20 amino acids of CDRH3 line the lateral side of the S2 apex (Fig. 4D) form four main stabilizing H-bonds. There, CDRH3 residues R102, Y105, S108, and Y111 form hydrogen bonds with N848, K733, N737, and R765 of the spike, respectively. CDRH3 R102 also forms a salt bridge with N848 in S2. The CDRH2 forms additional hydrogen bonds between N52 of the antibody and Q853 of spike, N54 with L849, S55, N745 and Q853, and Q58 of the antibody with E748 and Q853 of S2. Y60 in antibody FR2 forms an additional hydrogen bond with residue S750 of the spike. Light chain N92 and N93 interact with N737 and R765 of S2.

Access to the 6C10 epitope requires a dynamic opening of the spike. There are two polypeptide sections that contribute to the epitope: residues 733-772 and 844-862. Residues 733-772 are fully occluded in the closed, complete spike. Further, in the closed conformation of spike, the heavy chain of 6C10 would clash with the position of the S1 N-terminal domain of the neighboring closed protomer, and the light chain of 6C10 would clash with the S1 C-terminal portion adjacent to the furin cleavage site. Based on the structure, those clashes could not be resolved by mere positioning of the RBDs “up” or “down”, or by the natural motion of the NTD necessary for lifting of the RBD - the spike protomers must breathe open relative to each other. The second section of the epitope, residues 844-862 have become disordered. This region is accessible in the full-length spike and might be a starting point for, or sufficient for antibody binding. That hypothesis is supported by the fact that the two regions that form the 6C10 epitope are separated from each other in the postfusion conformation, although 6C10 can still recognize S2 in the postfusion conformation in addition to complete spike (Suppl Fig. S6).

Despite the apparent complexity of the conformational epitope recognized by 6C10, the antibody community S2-C (competition group) to which 6C10 belongs is abundant among the panel of anti-S2 antibodies presented in this study (Fig. 2B). The panel of mAbs was initially sorted using an S-2P full-length spike, which is pre-fusion, but has a demonstrated tendency to “breathe” (*30*). Moreover, residues contained in the footprint of 6C10 are highly conserved: 7 out of the 10 residues on the S2 that form H-bonds to 6C10 are conserved among the five human infective β-CoVs (Fig. 4E).

## DISCUSSION

Since the outbreak of COVID-19, numerous vaccines have been developed and several have been authorized for use in humans. However, the emergence of variants of concern bearing numerous mutations has evaded antibody responses, particularly against S1-containing epitopes (Suppl Fig. S7). Reactivity against S2 is broader and more durable against these variants, and may have played important roles in mitigating and preventing SARS-CoV-2 disease. Further, it has also been reported that pre-existing immunity against S2 epitopes conserved with common cold coronaviruses may have also mitigated SARS-CoV-2 severity (*17, 40–42*). Notably, the B cells of COVID-19 patients have increased somatic hypermutation and higher affinity against spike proteins of common cold coronaviruses (HCoV) than do pre-pandemic individuals, indicating that SARS-CoV-2 exposure might train the HCoV-induced B cells to produce more effective SARS-CoV-2 NAbs (*43–45*).

In this study, we present the structure-based design of a prefusion stabilized S2 ectodomain, with high thermostability, yield, and homogenous maintenance of a pre-fusion conformation. Five versions of S2 were engineered, which each yield a stable, prefusion molecule in the absence of an antibody or an S1 subunit. These stabilized S2 ectodomains in isolation did not yield a high-resolution map, but did yield low-resolution (∼7 Å) cryoEM maps. These maps and nsEM reconstructions confirm the prefusion shape of stabilized S2 in the absence of a bound antibody (Fig. 1C). Complexing the engineered S2-DS5 with an antibody isolated from a SARS-CoV-2 convalescent patient allowed a higher resolution, 2.9 Å cryo-EM structure to be obtained. The conformation of S2 is identical between the 7 Å unbound and the 2.9 Å antibody-bound structures.

### Conformational changes on the S2 and repositioning of the fusion peptide

It has been suggested that full spike undergoes a sequential motion where lifting of RBDs to the “up” position reduces the contacts of the S1 to S2 and allows S2 to “breathe” in a reversible motion from closed, to open and back (*30*). This “breathing” of the SARS-CoV-2 spike has been observed biophysically, and postulated by many (*46*) (*47*), but not yet visualized. All structures thus far for SARS-CoV-2 spike are of a closed conformation. For respiratory syncytial virus (RSV) and HIV, an open conformation has been visualized at very low resolution (*24, 27, 28*). Molecular simulations of ‘breathing and tilting’ have been also described for the flu HA, suggesting that these motions could potentially affect the attachment to the host cell and the release of progeny viruses, and the open conformation is accessible to a broadly protective human antibody, FluA-20, suggesting that keeping it in that position could be a way to develop more effective drugs against the flu virus (*48*). In this work, we present the first open conformation of a fusion subunit of a class 1 fusion protein at high resolution. We also reveal that adopting the open conformation is essential for full recognition by an antibody identified in convalescent individual, and likely others in its competition group.

We suggest that the opening motion of S2 might be driven by residues in the recently described fusion peptide (FP, residues 876-909) (*38*). Of these residues, Phe888 in the center of the fusion peptide might be particularly important. Phe888 is fully hidden from solvent in the closed conformation, but in the open conformation is the main residue responsible for the new interactions described here between the fusion peptide loop 885-897. Further, in the open conformation, Phe888 is fully solvent accessible. Note that Phe 888 is completely conserved among all human infective CoVs, and that in the subsequent, triggered, postfusion, six-helix bundle conformation, Phe888 is located in the center of the fusion peptide inserted into the target membrane (*38*). Short peptides designed from the internal fusion peptide region of SARS-CoV-2 S2 have shown high conformation-dependent inhibitory activity against the virus (*49*). Hence, taken together, this structure, of an open, breathing conformation of the FP may illuminate steps in the fusion cancade and provide a new template for fusion inhibition.

### A potential role for the open, breathing state in spike function and spike vaccination

Although it has not been directly demonstrated whether reversible breathing occurs in wild-type spikes on virions, it is possible that the ‘open’ S2 may also reflect an intermediate along the pathway to S1 shedding, during the transition from the prefusion conformation to the postfusion conformation. If the ‘open’ stage is an on-pathway intermediate, antibodies or ligands that trap the protein in this state may block the protein along the pathway to fusion.

6C10 recognizes two polypeptidic portions at the apex of S2: residues 733-772 and 844-862, with at least ten specific hydrogen bonds to 733-772 and at least six hydrogen bonds and a salt bridge to 844-862. Of these sections, 733-772 is fully masked in the closed spike, while 844-862 is accessible. One of these sections alone may be sufficient for binding, as 6C10 also recognizes post-fusion S2 in which 733-772 and 844-862 are too far apart to be simultaneously recognized by the same Fab fragment. Based on solvent accessibility, one might expect region 844-862 to be the initial attachment site for 6C10. Further, we believe initial attachment to 844-862 can best occur when S2 is in the pre-fusion conformation as 844-862 is structurally different between pre- and post-fusion conformations, and antibody access to residues 844-862 is sterically blocked in the post-fusion conformation Similarly, mice immunized with an S-2P version of MERS S induced antibodies with epitopes present only in the prefusion core of β-coronaviruses (980–1006) (*2, 21*), suggesting that these antibodies may have also been selected only by a spike in the open conformation.

Although the 6C10 epitope is visualized here at high-resolution in the context of an S2-only structure, we believe this epitope to exist and be available in a conformation also available to the complete spike in the prefusion conformation. Antibodies in the same competition group as 6C10 were also identified in individuals vaccinated with S-2P, a form of spike stabilized to favor the pre-fusion conformation (*50–52*), but not yet infected. 6C10 also recognizes VFLIP formats of SARS-CoV-2 and SARS-CoV-1 spike with additional proline substitutions to more completely eliminate pre- to post-fusion conformational transitions of spike (*33, 34, 53*), further suggesting that the epitope must be accessible in the presence of S1 and in the prefusion S1/S2 complex. The fact that 6C10 was identified in a person early in the pandemic, who was convalescent from COVID-19, but had not yet been vaccinated, suggests that these epitopes are exposed in natural infection, yet 6C10 doesn’t neutralize nor protect (Suppl Fig S5). Further, our studies on the array of S2-reactive antibodies, from both infected (but not yet vaccinated) and vaccinated (but not previously infected) individuals, demonstrate that such epitopes are present in the vaccine-elicited repertoire as well as the natural infection-elicited repertoire. We believe that breathing of the SARS-CoV-2 spike occurs in natural infection and in two-proline stabilized S-2P spike vaccination.

### The alternative conformation presents new druggable sites

Antibodies against conserved epitopes in S2 offer hope for recognition of variants and coronaviruses yet to emerge. In general, antibodies against S2 do not neutralize in cell culture as effectively as those against S1, although the capacity to neutralize may depend on how the assay is organized (presence or absence of TMPRSS2, different cell types, for example. Further, antibodies may be protective in the absence of neutralization by tagging infected cells for destruction, or through a mechanism not recapitulated in monotypic neutralization assays. Fewer antibodies against S2 may also be described thus far in the literature, because S2 is less immunogenic. Boosting with an S2-only subunit would improve elicitation of such antibodies, but was previously thwarted by the lack of a prefusion S2-only antigen in the pre-fusion conformation with which to achieve boosts. Antibodies against S2 may also be less prevalent due to the previous technical challenges with identifying and characterizing such antibodies, even if they had been present in vaccinees or convalescent individuals. The provision here of a prefusion S2-only protein will facilitate the identification and characterization of more such S2-reactive antibodies.

## ACKNOWLEDGEMENTS

We acknowledge National Institutes of Health grant NIAID P01 AI165072 and private philanthropic support (EOS, SS). We thank the cryoEM facility of La Jolla Institute for Immunology for data collection and assistance with processing of cryoEM data, and the John and Sue Major Center for Clinical Discovery at La Jolla Institute for Immunology for assistance with donor recruitment.

## AUTHOR CONTRIBUTIONS

Conceptualization, E.O., K.M.H. and E.O.S.; Methodology, E.O., R.R., K.M.H. and E.O.S.; Materials, E.O., R.R., F.A.S., M.A.Z., S.H.H., K.M.H.; Investigation E.O., R.R., R.D.A., S.H.H., K.M.H. Formal Analysis, E.O., R.R., R.D.A., D.Z., K.M.H. and E.O.S.; Writing – Original Draft, E.O. and E.O.S.; Writing – Review & Editing, E.O., K.M.H., E.O.S., Supervision, E.O.S.; Resources and Funding Acquisition, E.O.S.

## SUPPLEMENTARY LEGENDS

## STAR METHODS

## RESOURCE AVAILABILITY

### Lead Contact

Further information and requests for resources and reagents should be directed to and will be fulfilled by the Lead Contact, Erica Ollmann Saphire).

### Materials Availability

All reagents generated in this study are available from the Lead Contact upon request with a materials transfer agreement.

### Data and Code Availability

Datasets generated during this study are included in the article or are available from the corresponding authors on request. The cryo-EM map generated during this study is deposited in the Electron Microscopy Data Bank EMBL-EBI under accession code EMD-42985. The atomic model generated during this study is deposited in the RCSB Protein Data Bank. under accession code 8V5V.

## EXPERIMENTAL MODEL AND SUBJECT DETAILS

### Bacterial strains

*E. coli* strain Rosetta DE3 (Novagen) was grown in lysogeny broth. The genotype is: F-ompT hsdSB (rB-mB) gal dcm (DE3) pRARE (CamR). Selection markers were used at the indicated concentrations: kanamycin (100 μg/mL); chloramphenicol (28.3 μg/mL).

### Cell lines

Expi-CHO cells were obtained from Thermo Fisher Scientific and maintained in ExpiCHO-Expression Medium (Thermo Fisher Scientific). Vero CCL-81 cells were purchased from ATCC and maintained in Dulbecco’s Modified Eagle’s Medium (DMEM; Corning) supplemented with 2% FBS, 1% penicillin-streptomycin, 1% HEPES buffer, and 1% non-essential amino acids

## METHOD DETAILS

### Design of SARS-CoV-2 S2 variants

SARS-CoV-2 S2 variants (residues 691-1211) were initially designed using the VFLIP construct Wuhan strain (Genbank: MN908947), which contains 5 proline substitutions (F817P, A892P, A899P, A942P, and V967P), and an inter-protomer disulfide bond formed by Y707C-T883C. All variants were cloned into a PhCMV mammalian expression vector containing a C-terminal foldon trimerization domain followed by an HRV-3C cleavage site and a Twin-Strep-Tag.

### Transient transfection and protein purification

SARS-CoV-2 S2 variants were transiently transfected in ExpiCHO-S cells (Thermo Fisher). CHO cells were maintained and transfected according to the manufacturer’s protocols. For all ExpiCHO cultures, the manufacturer’s “High Titer” protocol was used with a 7-day culture incubation to assess relative expression. Briefly, plasmid DNA and Expifectamine were mixed in Opti-PRO SFM (Gibco) according to the manufacturer’s instructions and added to the cells. On day 1, cells were fed with manufacturer-supplied feed and enhancer as specified in the manufacturer’s protocol, and cultures were moved to a shaker incubator set to 32 °C, 5% CO_2_ and 115 RPM. On day 7, cultures were clarified by centrifugation, followed by addition of BioLock (IBA Life Sciences), passage through a 0.22 µM sterile filter, and purification on an DKTA go system (Cytiva) using a 5mL Strep-Tactin XT column equilibrated with TBS buffer (25mM Tris pH 7.6, 200mM NaCl, 0.02% NaN3), and eluted in TBS buffer supplemented with 10mM d-desthiobiotin (Sigma Aldrich). Proteins were then purified by size-exclusion-chromatography (SEC) on a Superdex 6 Increase 10/300 column (Cytiva) in the same TBS buffer.

### Differential Scanning Calorimetry

Thermal stability of the SARS-CoV-2 S2 constructs was analyzed by Nano DSC (TA instruments). The corresponding protein (500 µg) was buffer-exchanged to phosphate-buffered saline (PBS) and loaded into the sample wells. The temperature ramped from 20 °C to 100°C at 1 °C per minute. The resulting thermogram was corrected by subtracting a buffer blank data set and baseline-correcting before fitting to a thermodynamic model to extract the exact melting temperature (Tm).

### Negative stain electron microscopy (NS-EM)

NS-EM was utilized to visualize the SARS-CoV-2 S2 designs, as well as the complexes formed with the S2-DS5 and the Fabs fragments of the human mAbs 6C10, 1A11, 6E1, and 4B3. To prepare the complexes, S2-DS5 and Fab in a 1:2 molar ratio, were incubated overnight at 4D. The samples were then injected over a gel filtration column (Superose 6 10/30, GE Life Sciences) equilibrated with 20mM Tris pH 8.0 and 150mM NaCl. The complex peak was diluted to 0.02mg/ml, and 3 µL of the peak was applied to a carbon film 400 mesh grid for 1 minute. The grid was then washed three times with 20 µL Milli-Q water and stained three times with 20 µL drops of 1% uranyl formate, with the first two drops briefly, and the third time for 1 minute. The grid was blotted with Whatman filter paper after each application of liquid and air dried. Micrographs were collected on a Titan Halo, operating with an accelerating voltage of 300kV and fitted with a Gatan K3 camera. We used a magnification corresponding to a pixel size of 1.7A/pixel.

### Beacon-based antibody discovery

Activated memory B cells were loaded onto OptoSelect 11K chips (Bruker) and isolated as single cells in nanoliter pens using OEP light cages. Cells were screened in a 30 min time course assay for secretion of antibodies that bound to streptavidin beads (Spherotech) coated with 10µg/mL biotinylated SARS-CoV-2 S-2P, as reported previously (*39*). Secreted antibodies were detected with 1µg/mL goat anti-human IgG (H+L)-Alexa Fluor 594 (Invitrogen), which was added to the cell culture media used to resuspend the antigen-coated beads.

Synthesis of cDNA from antigen-positive cells was carried out on-chip using the OptoSeq BCR kit (Bruker), according to the manufacturer’s directions. First-strand reaction products were exported on mRNA capture beads and deposited into individual wells of a 96-well plate. Total cDNA was amplified using Platinum SuperFi II polymerase (Invitrogen). After enzymatic cleanup, antibody heavy and light chain variable domains were amplified with two rounds of nested PCR using Platinum II Hot-Start polymerase (Invitrogen). The resulting PCR products were assessed using 96w E-gels (Thermo Fisher) and paired wells were Sanger sequenced. Sequences were annotated using the bioinformatics platform PipeBio (PipeBio ApS, Horsens, DK).

Unique VH and VL domains were cloned into linearized human antibody expression vectors (human IgG1 and kappa light chain) using Gibson assembly (NEB) according to the manufacturer’s directions. Ligation reactions were transformed into 5-alpha F’Iq competent E. coli cells (NEB). QIAprep 96 Turbo Miniprep kits (Qiagen) were used to isolate plasmids according to the manufacturer’s instructions. Briefly, S block wells containing 1.1 mL LB media spiked with antibiotic were inoculated with single colonies and incubated overnight at 37 °C with agitation. DNA extraction was carried out with Qiagen buffer solutions and protocols. Plasmids were sequenced to ensure that the genes were in-frame, and that the cloned heavy and light chain variable domains matched PCR sequences.

### High-throughput SPR binding kinetics

Binding kinetics measurements were performed on the Carterra LSA platform using a CDMP sensor chip (Carterra). The chip was activated with a freshly prepared solution of 130 mM 1-ethyl-3-(3-dimethylaminopropyl) carbodiimide hydrochloride (EDC) (Pierce PG82079) and 33 mM N-hydroxysulfosuccinimide (Sulfo-NHS) (ThermoFisher Scientific 24510) in 0.1 M MES pH 5.5 using the SFC. A surface capture lawn was prepared with 50 ug/mL of goat anti-Human IgG Fc secondary antibody (VWR, 103255-066) in 10 mM sodium acetate (pH 4.5)/0.01% Tween. Unreactive esters were quenched with a 7-minute injection of 1 M ethanolamine-HCl (pH 8.5). Purified mAbs were captured in quadruplicate using the 96PH with 1X HBSTE buffer (10 mM HEPES pH 7.4, 150 mM NaCl, 3 mM EDTA and 0.01% Tween-20) as running buffer and antibody diluent.

A two-fold dilution series of the spike construct shown in Figure 2 was prepared in 1xHBSTE-BSA buffer (10 mM HEPES pH 7.4, 150 mM NaCl, 3 mM EDTA, 0.05% Tween-20, supplemented with 0.5 mg/ml BSA). Protein was injected onto the chip surface (at 25 °C) using the SFC, from the lowest to the highest concentration, without regeneration in between. Five buffer injections of the buffer before the lowest non-zero concentration were used for signal stabilization. For each concentration, baseline data were collected for 120 seconds, association data for 300 seconds and dissociation data for 900 seconds. After the titration of each analyte, the huFc capture lawn surface was regenerated with two pulses (17 seconds per pulse) of 10mM Glycine, pH 2.0 and mAbs re-captured in between each analyte. The running buffer for all kinetic steps was 1xHBSTE-BSA.

Titration data were processed with the Kinetics software package (Carterra), including reference subtraction, buffer subtraction and data smoothing. Analyte binding time courses for each antibody were fitted to a 1:1 Langmuir model to derive ka, kd and K_D_ values.

### High-throughput SPR epitope binning

Epitope binning was performed on the Carterra LSA^®^ HT-SPR using an HC30M sensor chip (Carterra). The chip was activated with a freshly prepared solution of 130 mM 1-ethyl-3-(3-dimethylaminopropyl)carbodiimide hydrochloride (EDC) (Pierce PG82079) and 33 mM N-hydroxysulfosuccinimide (Sulfo-NHS) (ThermoFisher Scientific 24510) in 0.1 M MES pH 5.5 using the SFC. Antibodies were diluted to 5 µg/mL in 10 mM sodium acetate (pH 4.5) /0.01% Tween and immobilized in quadruplicate using the 96PH for 15 minutes. Unreactive esters were quenched with a 7-minute injection of 1 M ethanolamine-HCl (pH 8.5).

Epitope binning for S2-DS5 was performed with a premix assay format at 25 °C. A mixture of 50 nM of S2-DS5 was preincubated with 200 nM of each analyte antibody for at least 30 minutes before injecting over the array for 4 minutes. The sample injection for every sixth cycle was S2-DS5 only instead of antibody-S2-DS5 mixture. The surface was regenerated each cycle with double pulses (30 seconds per pulse) of 10 mM Glycine pH 2.0. Data was processed and analyzed with Epitope® software (Carterra). Data was referenced using unprinted locations (nearest reference spots) on the array for both experimental set ups. The response of each injection cycle was normalized to the response level in S2-DS5 only cycles. Competition results were visualized as a heat map that depicts blocking relationships of analyte/ligand pairs. Clones that suffered from severe loss of activity or lack of complete dissociation from S2-DS5 when used as ligands were excluded from analysis. Antibodies with similar patterns of competition are clustered together in a dendrogram and are assigned to shared communities.

### Antibody expression

25ml culture of ExpiCHO cells were maintained as described above. On the day of transfection, plasmids encoding heavy and light chains (10 µg each) were mixed with 80 µl Expifectamine, allowed to sit for 1-2 minutes, and added to cells dropwise over 5 minutes. The flask containing the transfected cells was allowed to shake at 120 rpm in an incubator maintained at 37 °C, 8% CO_2_. The cells were fed 18-22 hours after transfection with a mixture of 6 ml cold feed and 150 µl cold enhancer and moved to a 32 °C incubator with 5% CO_2_ and incubated with agitation at 120 rpm.

### Antibody purification

The cells were harvested 7-8 days post transfection by centrifugation at 4,000 g for 30 minutes. The supernatant was filtered using a 0.22 µm filter and the pH was adjusted to 7.5. Packed protein A beads (2 ml; #Praesto AP Purolite Life Sciences, PR00300-310) were washed with TBS and incubated with the supernatant for between 30 min and 2 hrs at RT. The mixture was passed through a gravity column and the beads were washed with 20 CV TBS to remove unbound proteins; IgG was eluted using 6 ml of elution buffer (100 mM glycine pH 2.2). Neutralization buffer (900 µl, 1M Tris pH 8) was added to bring the pH to neutral. IgG was dialyzed into TBS pH 7.5 overnight at 4 °C. Aliquots (1mg/ml) were made and stored at -20 °C.

### Digestion of IgG to Fabs

Each antibody IgG (2mg) was incubated with 4% papain w/w for 2 hours at 37 °C. L-cysteine (Sigma) was added to a final concentration of 15 mM and incubated for 1 hour at 37 °C. Digestion was quenched with 50 mM iodoacetamide. The Fc portion was removed by passing the mixture over a column containing protein A beads. Fab was recovered from the flowthrough and buffer-exchanged to PBS using a Vivaspin 10k concentrator.

### Cryo-EM data collection and processing

The S2-DS5/6C10_Fab complex was concentrated to 1 mg/ml and electron microscopy grids were prepared by placing a 3 µL aliquot of the sample on a plasma-cleaned C-flat grid (2/1C-3T, Protochips Inc) that was then immersed in liquid ethane for vitrification (VitroBot). Grids were loaded into a Titan Krios G3 electron microscope (Thermo Fisher Scientific) equipped with a K3 direct electron detector (Gatan, Inc.) at the end of a BioQuantum energy filter, using an energy slit of 20eV. The microscope was operated with an accelerating voltage of 300kV. Grids were imaged with a pixel size of 0.66 Å in counting mode. Data was acquired using the software EPU. Two independent batches of the complex S2-DS5/6C10_Fab were frozen. A total of 5022 and 7422 movie stacks were motion-corrected using the patch motion correction, and the CTF parameters were determined using the patch CTF estimation in Warp (*54*). Warp picker was used to select a total of 1,137,885 particles. After two-dimensional classification and several rounds of hetero-refinement and non-uniform refinement in cryoSPARC-v2 (*55*) (Structura Biotechnology Inc.), a set of 407,314 refinements yielded a 2.9 Å Cryo-EM density map with C1 symmetry (Supp. Fig S2).

### Cryo-EM model building and structure analysis

Structure prediction of the 6C10 Fab was performed with AlphaFold server (*56*). Model building was performed in Coot 0.9.8.7 ((*57, 58*) and guided by the PDB model 6XKL. Model refinement and validation was performed in Phenix 1.20 (*59*). For visualization purposes, the map was processed by DeepEMhancer (*60*) and ChimeraX 1.6 (*61, 62*) was used to prepare representations of the structure. The surface areas and interactions S2-DS5/6C10_Fav were analyzed using the PDBePISA server (*63*).

