## Supplementary figures for "Structure of a SARS-CoV-2 spike S2 subunit in a pre-fusion, open conformation"

### Slide 1
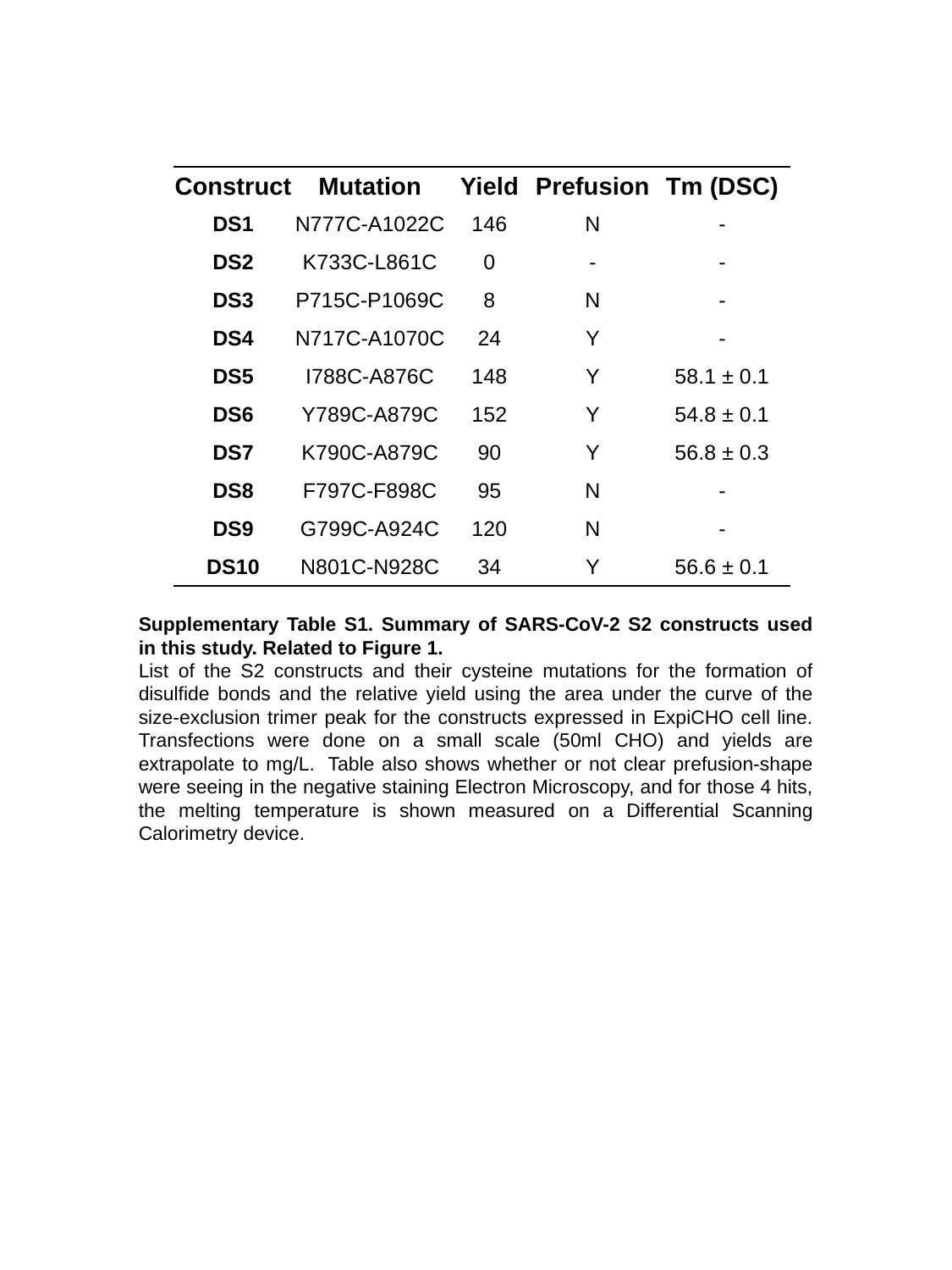

| Construct | Mutation | Yield | Prefusion | Tm (DSC) |
| --- | --- | --- | --- | --- |
| DS1 | N777C-A1022C | 146 | N | - |
| DS2 | K733C-L861C | 0 | - | - |
| DS3 | P715C-P1069C | 8 | N | - |
| DS4 | N717C-A1070C | 24 | Y | - |
| DS5 | I788C-A876C | 148 | Y | 58.1 ± 0.1 |
| DS6 | Y789C-A879C | 152 | Y | 54.8 ± 0.1 |
| DS7 | K790C-A879C | 90 | Y | 56.8 ± 0.3 |
| DS8 | F797C-F898C | 95 | N | - |
| DS9 | G799C-A924C | 120 | N | - |
| DS10 | N801C-N928C | 34 | Y | 56.6 ± 0.1 |
Supplementary Table S1. Summary of SARS-CoV-2 S2 constructs used in this study. Related to Figure 1.
List of the S2 constructs and their cysteine mutations for the formation of disulfide bonds and the relative yield using the area under the curve of the size-exclusion trimer peak for the constructs expressed in ExpiCHO cell line. Transfections were done on a small scale (50ml CHO) and yields are extrapolate to mg/L.  Table also shows whether or not clear prefusion-shape were seeing in the negative staining Electron Microscopy, and for those 4 hits, the melting temperature is shown measured on a Differential Scanning Calorimetry device.

### Slide 2
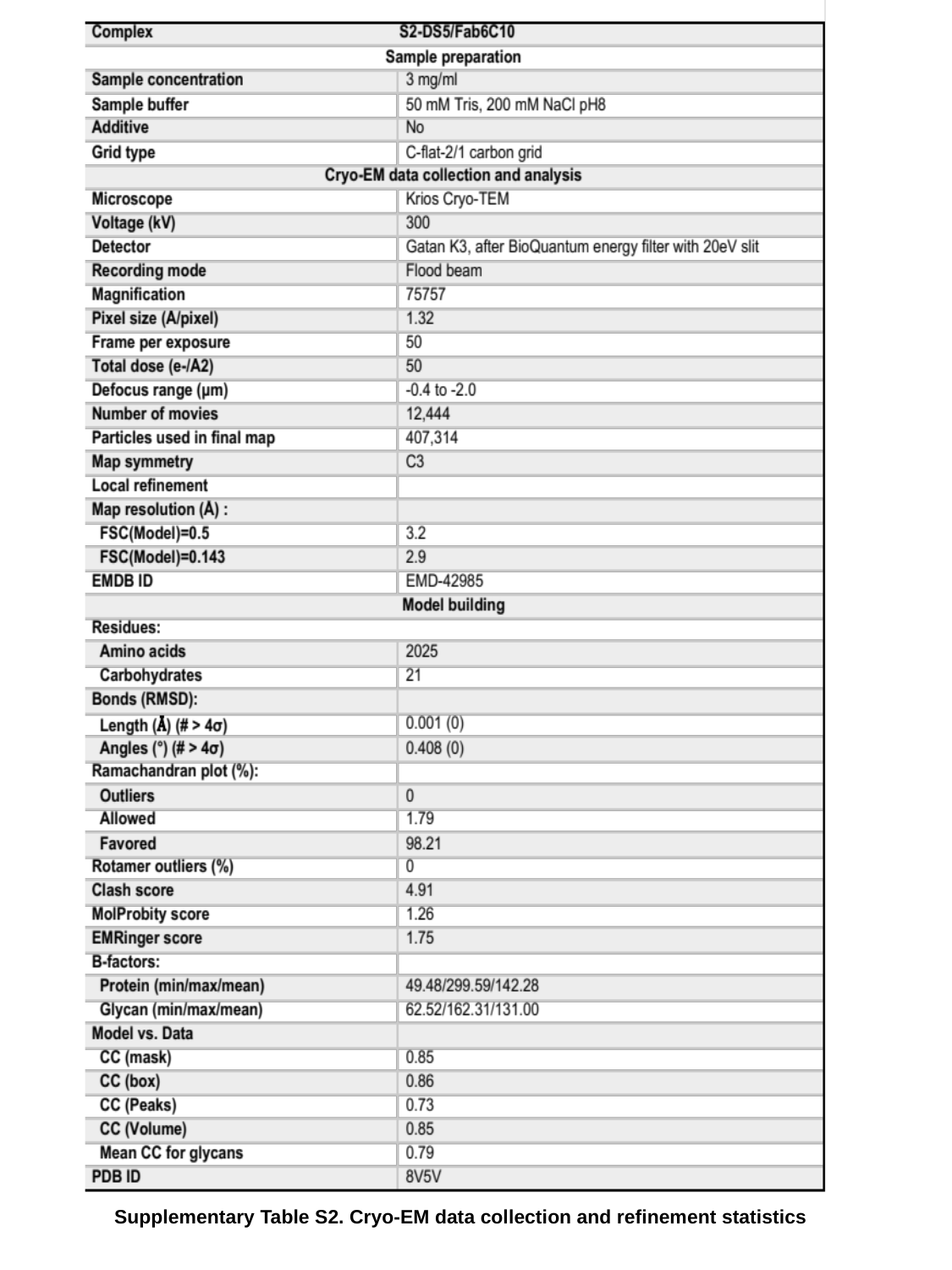

Supplementary Table S2. Cryo-EM data collection and refinement statistics

### Slide 3
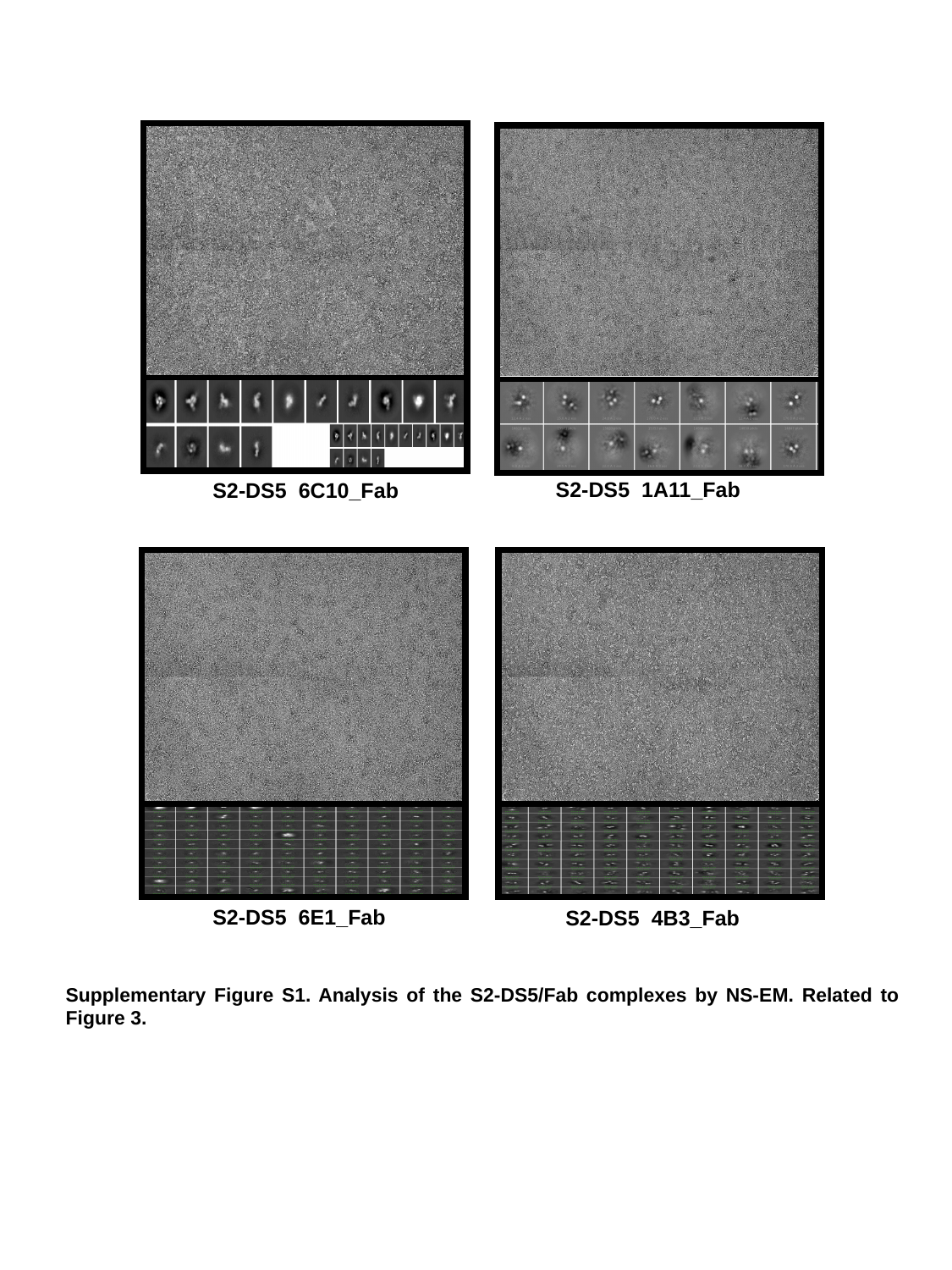

S2-DS5 6C10_Fab
S2-DS5 1A11_Fab
S2-DS5 6E1_Fab
S2-DS5 4B3_Fab
Supplementary Figure S1. Analysis of the S2-DS5/Fab complexes by NS-EM. Related to Figure 3.

### Slide 4
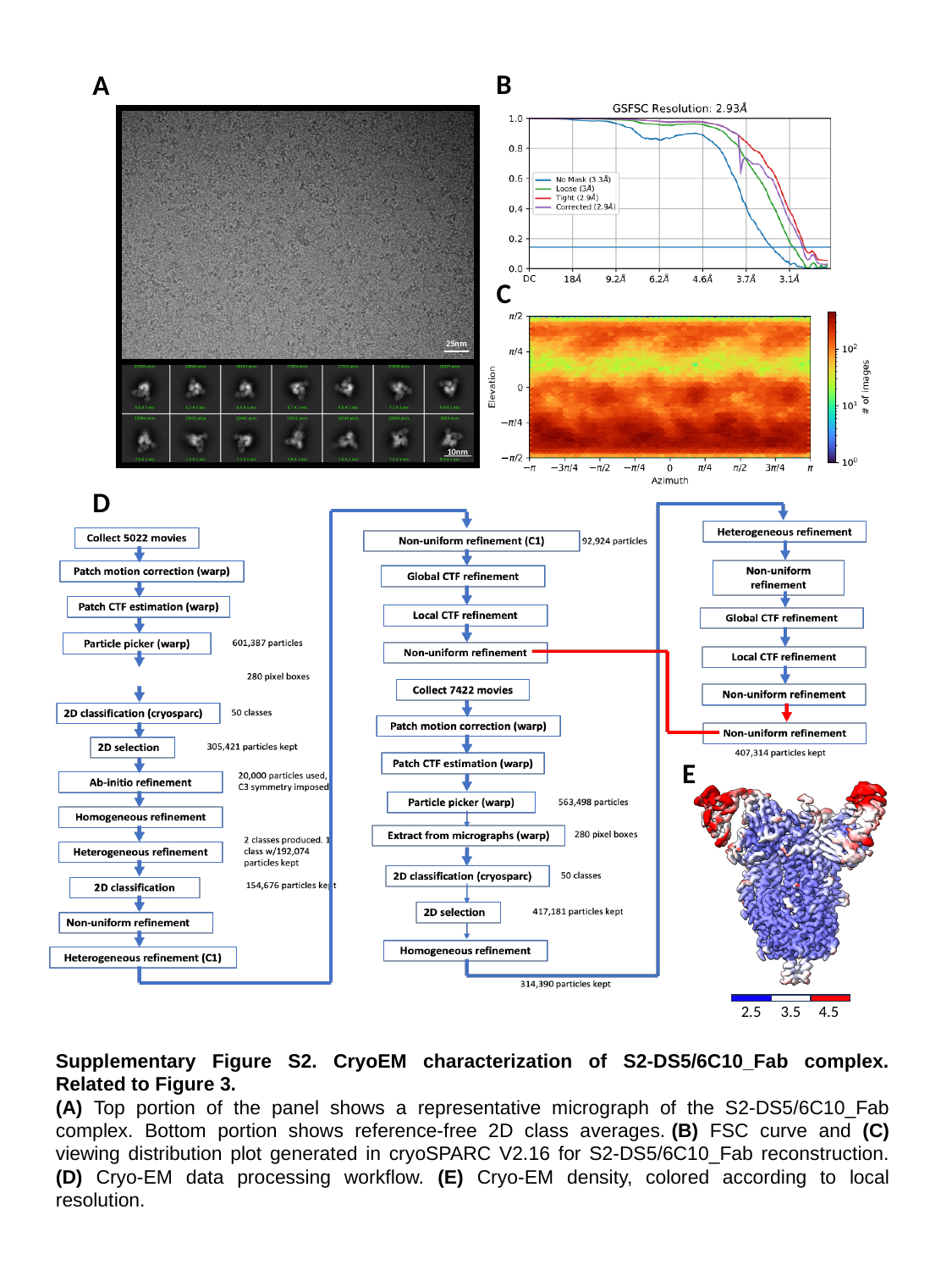

B
A
C
25nm
10nm
D
E
2.5
3.5
4.5
Supplementary Figure S2. CryoEM characterization of S2-DS5/6C10_Fab complex. Related to Figure 3.
(A) Top portion of the panel shows a representative micrograph of the S2-DS5/6C10_Fab complex. Bottom portion shows reference-free 2D class averages. (B) FSC curve and (C) viewing distribution plot generated in cryoSPARC V2.16 for S2-DS5/6C10_Fab reconstruction. (D) Cryo-EM data processing workflow. (E) Cryo-EM density, colored according to local resolution.

### Slide 5
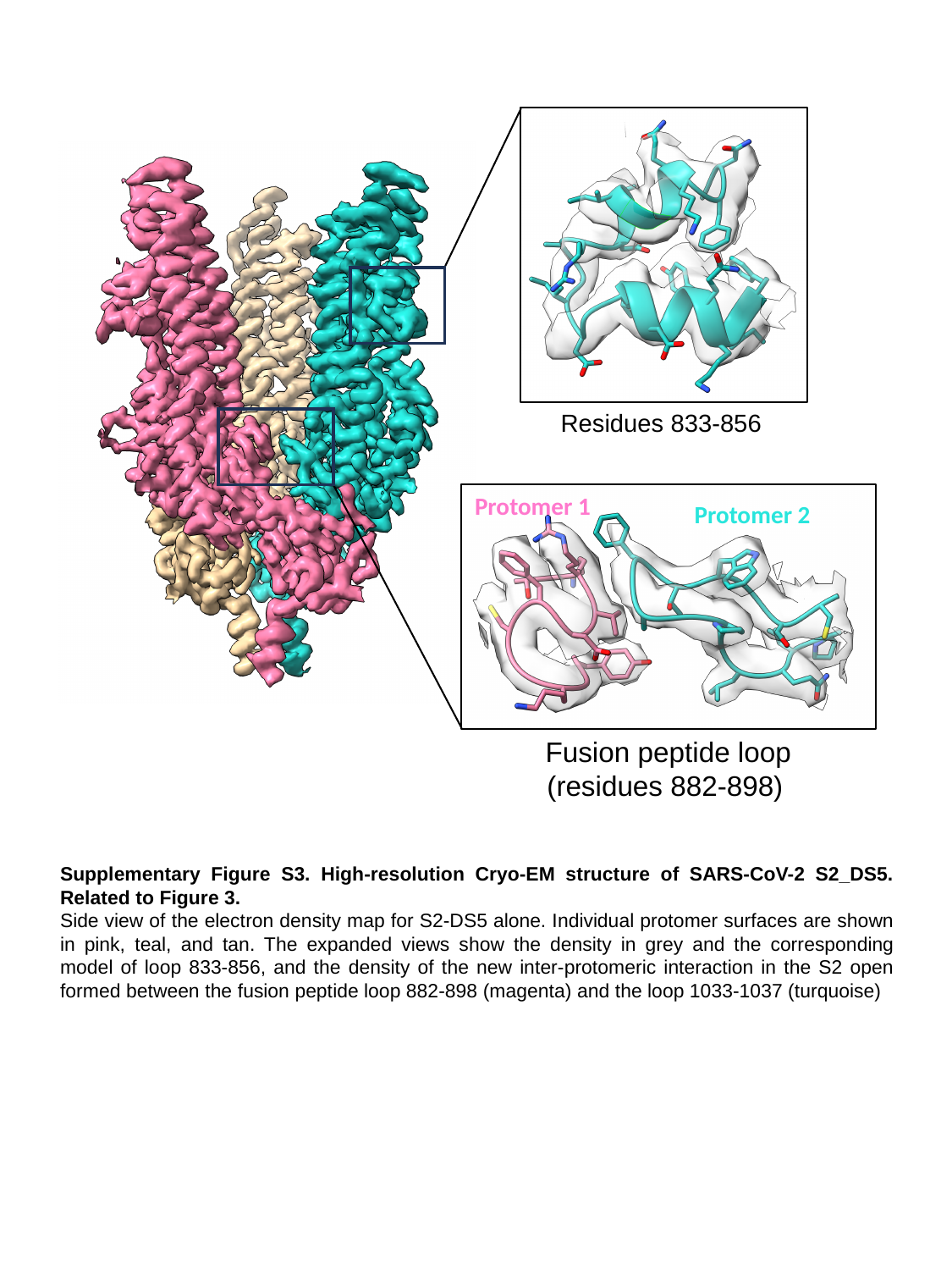

Residues 833-856
Protomer 1
Protomer 2
Fusion peptide loop (residues 882-898)
Supplementary Figure S3. High-resolution Cryo-EM structure of SARS-CoV-2 S2_DS5. Related to Figure 3.
Side view of the electron density map for S2-DS5 alone. Individual protomer surfaces are shown in pink, teal, and tan. The expanded views show the density in grey and the corresponding model of loop 833-856, and the density of the new inter-protomeric interaction in the S2 open formed between the fusion peptide loop 882-898 (magenta) and the loop 1033-1037 (turquoise)

### Slide 6
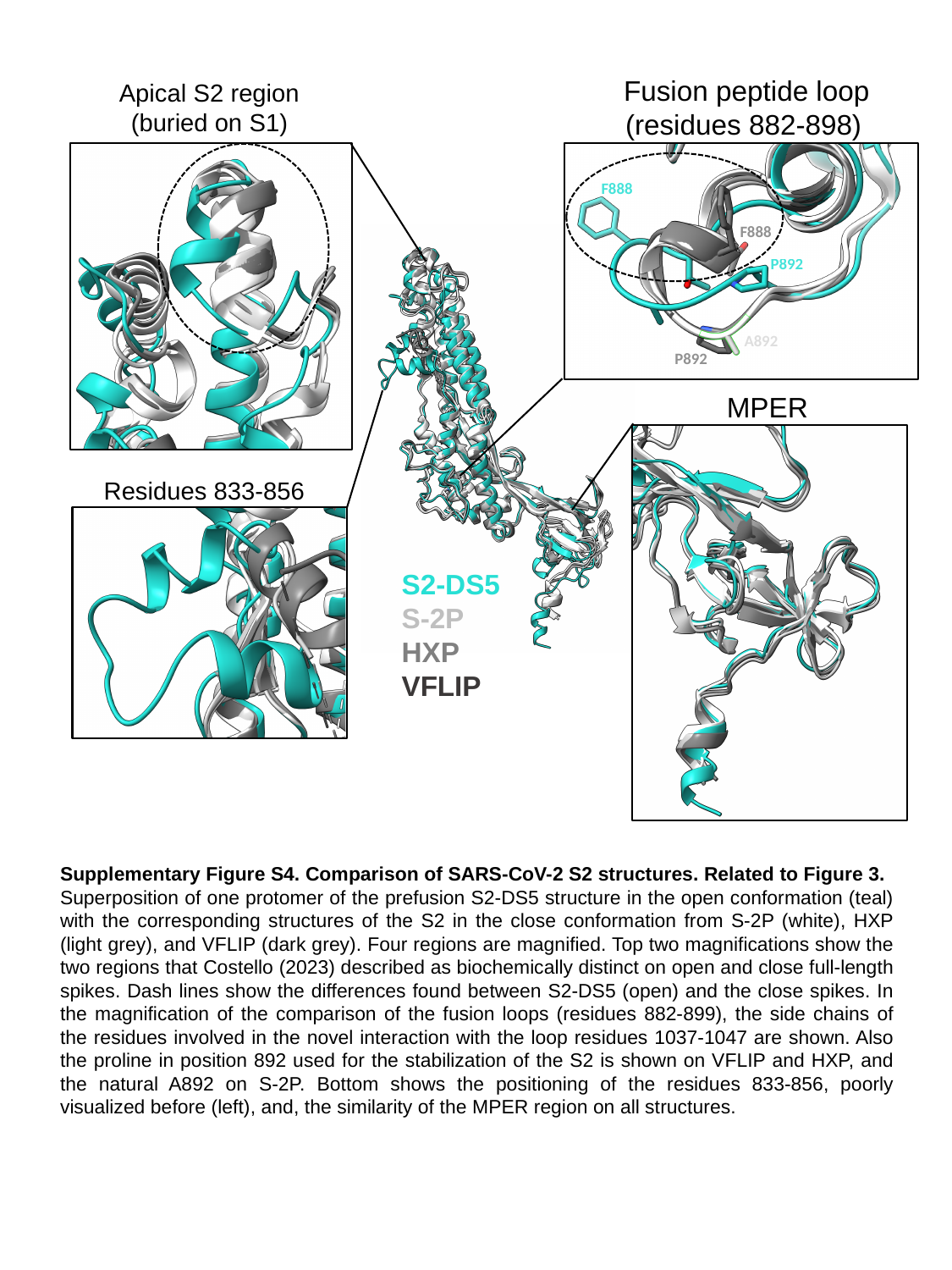

Fusion peptide loop (residues 882-898)
Apical S2 region
(buried on S1)
F888
F888
P892
A892
P892
MPER
Residues 833-856
S2-DS5
S-2P
HXP
VFLIP
Supplementary Figure S4. Comparison of SARS-CoV-2 S2 structures. Related to Figure 3.
Superposition of one protomer of the prefusion S2-DS5 structure in the open conformation (teal) with the corresponding structures of the S2 in the close conformation from S-2P (white), HXP (light grey), and VFLIP (dark grey). Four regions are magnified. Top two magnifications show the two regions that Costello (2023) described as biochemically distinct on open and close full-length spikes. Dash lines show the differences found between S2-DS5 (open) and the close spikes. In the magnification of the comparison of the fusion loops (residues 882-899), the side chains of the residues involved in the novel interaction with the loop residues 1037-1047 are shown. Also the proline in position 892 used for the stabilization of the S2 is shown on VFLIP and HXP, and the natural A892 on S-2P. Bottom shows the positioning of the residues 833-856, poorly visualized before (left), and, the similarity of the MPER region on all structures.

### Slide 7
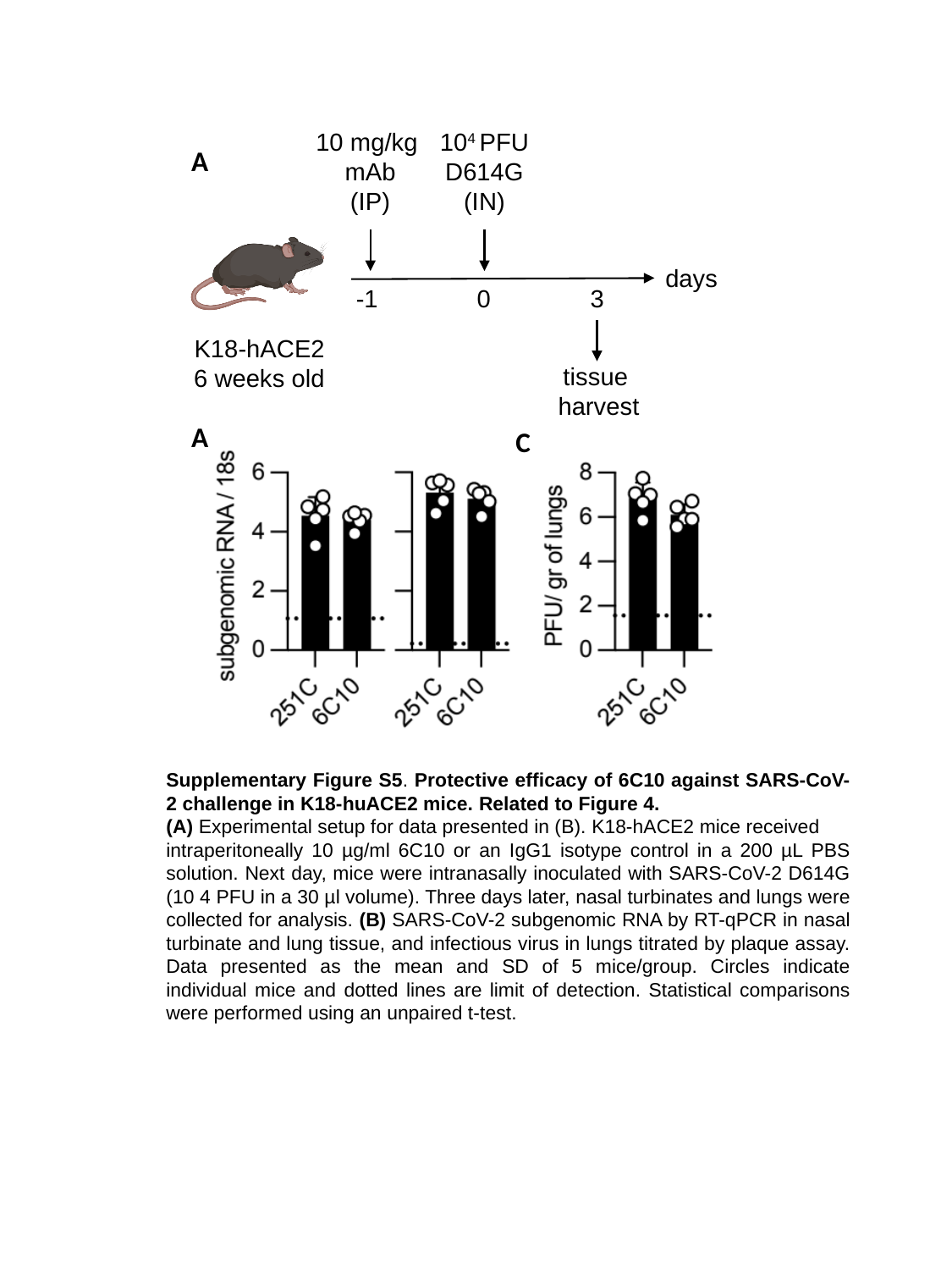

10 mg/kg
mAb
(IP)
10^4^ PFU
D614G
(IN)
A
days
-1
0
3
K18-hACE2
6 weeks old
tissue
harvest
A
C
Supplementary Figure S5. Protective efficacy of 6C10 against SARS-CoV-2 challenge in K18-huACE2 mice. Related to Figure 4.
(A) Experimental setup for data presented in (B). K18-hACE2 mice received
intraperitoneally 10 µg/ml 6C10 or an IgG1 isotype control in a 200 µL PBS solution. Next day, mice were intranasally inoculated with SARS-CoV-2 D614G (10 4 PFU in a 30 µl volume). Three days later, nasal turbinates and lungs were collected for analysis. (B) SARS-CoV-2 subgenomic RNA by RT-qPCR in nasal turbinate and lung tissue, and infectious virus in lungs titrated by plaque assay. Data presented as the mean and SD of 5 mice/group. Circles indicate individual mice and dotted lines are limit of detection. Statistical comparisons were performed using an unpaired t-test.

### Slide 8
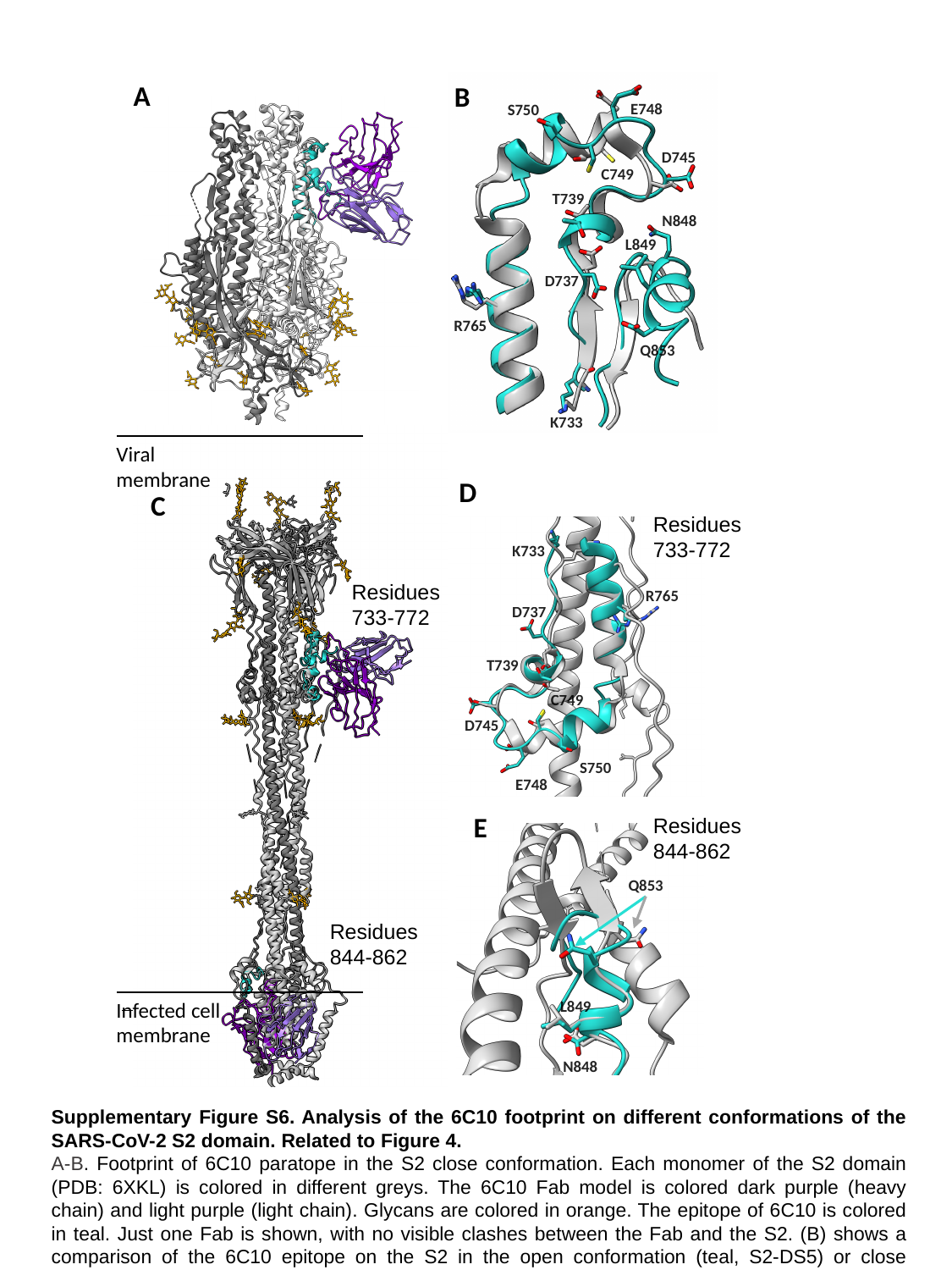

A
E748
S750
D745
C749
T739
D737
K733
B
N848
L849
R765
R765
Q853
Infected cell membrane
Viral membrane
D
C
Residues 733-772
K733
R765
D737
T739
C749
D745
S750
E748
Residues 733-772
E
Residues 844-862
Q853
L849
N848
Residues 844-862
Supplementary Figure S6. Analysis of the 6C10 footprint on different conformations of the SARS-CoV-2 S2 domain. Related to Figure 4.
A-B. Footprint of 6C10 paratope in the S2 close conformation. Each monomer of the S2 domain (PDB: 6XKL) is colored in different greys. The 6C10 Fab model is colored dark purple (heavy chain) and light purple (light chain). Glycans are colored in orange. The epitope of 6C10 is colored in teal. Just one Fab is shown, with no visible clashes between the Fab and the S2. (B) shows a comparison of the 6C10 epitope on the S2 in the open conformation (teal, S2-DS5) or close conformation (light gray). Side chains of the residues forming H-bonds with the Fab are shown in sticks. C-D. Footprint of 6C10 paratope in the S2 postfusion (PDB: 6FDW). (C) In the postfusion conformation, the 6C10 epitope is split, and the accessibility of 6C10 Fab is shown with two Fabs. (D-E) show a comparison of the 6C10 epitope on the S2 in the postfusion conformation of residues 733-772 (D) and 844-862 (E).

### Slide 9
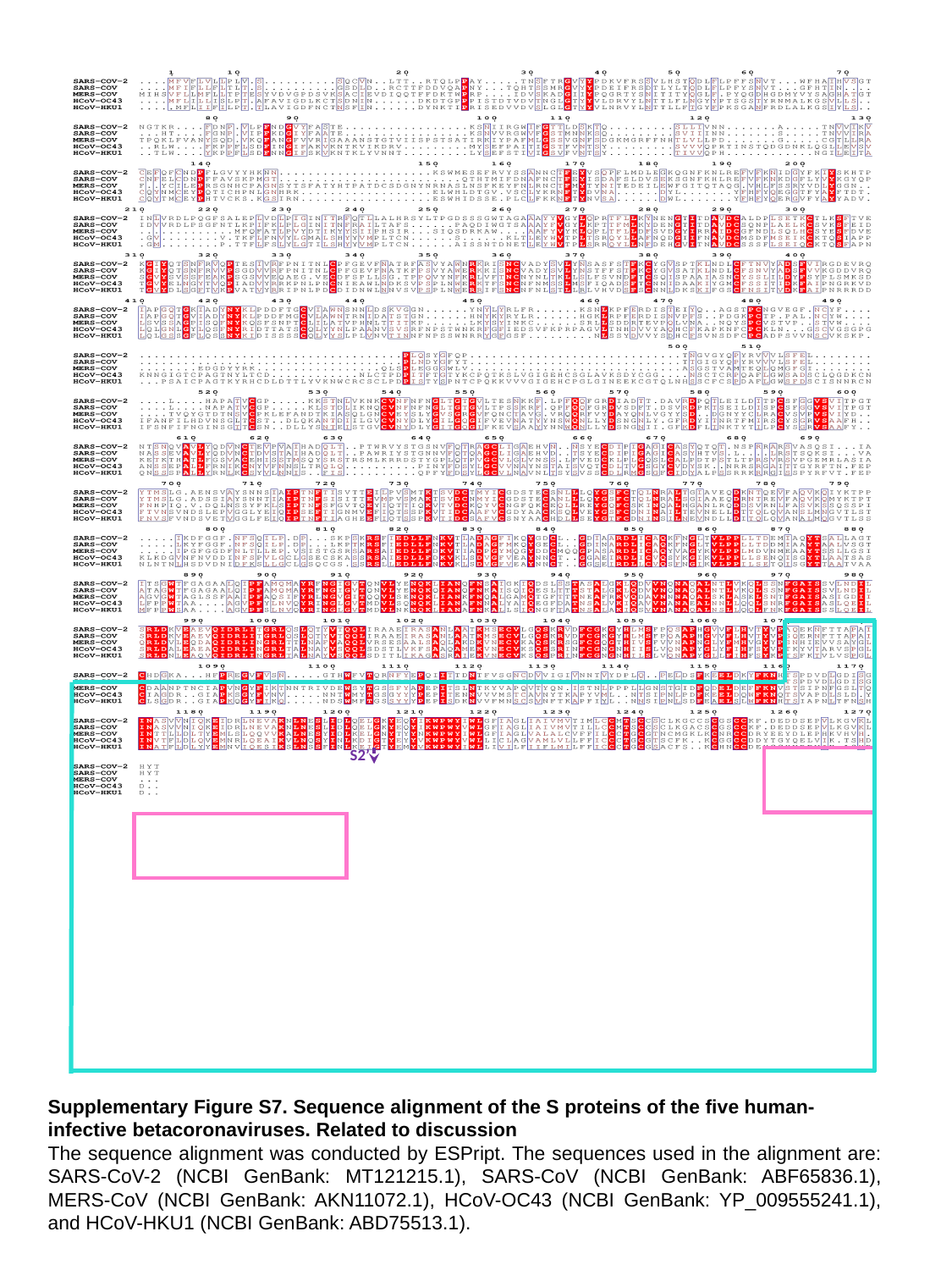

S2’
Supplementary Figure S7. Sequence alignment of the S proteins of the five human-infective betacoronaviruses. Related to discussion
The sequence alignment was conducted by ESPript. The sequences used in the alignment are: SARS-CoV-2 (NCBI GenBank: MT121215.1), SARS-CoV (NCBI GenBank: ABF65836.1), MERS-CoV (NCBI GenBank: AKN11072.1), HCoV-OC43 (NCBI GenBank: YP_009555241.1), and HCoV-HKU1 (NCBI GenBank: ABD75513.1).
